# *Allium roseum* L. extract inhibits amyloid beta aggregation and toxicity involved in Alzheimer’s disease

**DOI:** 10.1101/789651

**Authors:** Abdelbasset Boubakri, Manuela Leri, Monica Bucciantini, Hanen Najjaa, Abdelkarim Ben Arfa, Massimo Stefani, Mohamed Neffati

## Abstract

*Allium roseum* is an important medicinal and aromatic plant, specific to the North African flora and a rich source of important nutrients and bioactive molecules including flavonoids and organosulfur compounds whose biological activities and pharmacological properties are well known. In the present study, the inhibition of amyloid beta protein toxicity by the ethanolic extract of this plant is investigated for the first time. Preliminary biochemical analyses identified kæmpferol and Luteolin-7-o-glucoside as the more abundant phenolic compounds. The effects of *A. roseum* extract (ARE) on amyloid beta-42 (Aβ_42_) aggregation and aggregate cytotoxicity, were investigated by biophysical (ThT assay, Dynamic light scattering and transmission electron microscopy) and cellular assays (cytotoxicity, aggregate immunolocalization, ROS measurement and intracellular Ca^2+^ imaging). The biophysical data suggest that ARE affects the structure of Aβ_42_ peptide, inhibits its polymerization, and interferes with the path of fibrillogenesis. The data with cultured cells shows that ARE reduces Aß_42_ aggregate toxicity by inhibiting aggregate binding to the cell membrane and by decreasing both oxidative stress and intracellular Ca^2+^. Accordingly, ARE could act as a neuroprotective factor against Aβ aggregate toxicity in Alzheimer’s disease.

## 1. Introduction

Alzheimer’s disease (AD), the main cause of senile dementia, is a progressive neurodegenerative disorder associated with cognitive impairment and loss of neuronal cells affecting over 25 million people worldwide [1], thus representing a heavy economic burden and a major health problem. The pathology is characterized by the accumulation in the brain of amyloid-β peptide (Aβ) and hyperphosphorylated tau protein mainly found as extracellular senile plaques and intracellular neurofibrillary tangles, respectively [2]. Aβ oligomers overproduction caused by mutations in APP and presenilins [3, 4] appears to be directly associated to cognitive dysfunction and neurodegeneration in AD [5]. Accordingly, toxic Aβ oligomers production and/or interaction with nerve cells appear useful targets to reduce neuronal cell dysfunction. Feart, Samieri [6] suggest that in most cases AD is affected by a combination of genetic and environmental risk factors, including physical activity and nutrition; in particular, increasing evidence indicates that diet plays an important role in AD. Several studies have associated plant food (i.e., vegetables, fruits, legumes, and cereals) consumption to reduced risk of AD [7, 8]. Moreover, traditional medicine is increasingly appreciated as a useful approach to treat many illnesses, and large funds are invested to explore the therapeutic potential of medicinal plants. Fortunately, the search of active compounds from nature prospers and the results increasingly obtained confirm the need to better known the therapeutic potential of these natural drugstores.

Phenolic compounds, a very large class of plant molecules and among the most abundant bioactive substances in plant kingdom, have attracted the attention of many researchers for their therapeutic properties. They are known since long time for their antioxidant potential, being able to reduce, or even suppress, the oxidative stress in cells exposed to reactive oxygen species or heavy metals [9]. Recently, many studies showed that polyphenols induce neuroprotection and have beneficial effects against ageing and age-related pathological conditions [10] possibly including Alzheimer disease [11, 12]. In particular, these molecules exhibit a cytoprotective effect against Aβ aggregate toxicity to neuronal cells, both *in vivo* and *in vitro* [11, 12]. In a previous review Stefani and Rigacci [13] the neuroprotective power of natural phenols against Aβ toxicity was associated to their anti-amyloidogenic and anti-inflammatory activity, to the inhibition of production of toxic amyloid aggregates, to the protection against oxidative stress and to the activation of autophagy.

The use of herbal medicines was widely practiced in Tunisia. Among the different medicinal plants of the Tunisian flora, *Allium roseum* Var. *odoratissimum*, highly consumed by locals especially in the south of the Tunisia, is a rich source of organosulfur compounds and flavonoids, especially kaempherol [14]. Recent research showed that the administration of this flavonoid attenuates the oxidative stress caused by Aβ amyloids [15]. Furthermore, *A. roseum* is known for its therapeutic properties, particularly its antioxidant power [14, 16]. However, up until now there is a substantial lack of information about a possible use of this medicinal plant in the treatment or prevention of neurodegenerative diseases. The present study is the first investigation aimed at describing the effect of this medicinal plant on amyloid aggregation and toxicity of the Aβ_42_ amyloids involved in AD.

## 2. Material and Methods

### 2.1. Plant material and extract preparation

The whole plants of wild-growing *A. roseum* were collected from the arid South-East of Tunisia (Bengardane), at the vegetative stage of the plant cycle. The plant material was botanically authenticated according to the “Flora of Tunisia” [17]. Plant leaves were lyophilized and grinded to get a fine powder, and 5.0 g of the powder was extracted with 50 mL of absolute ethanol.

### 2.2. Concentrations of phenolic compounds

#### 2.2.1. Total Phenol Content (TPC)

Total phenol concentration of the *A. roseum* aqueous extracts was determined using the Folin-Ciocalteu reagent [18]. Briefly, 125 μL of extract were mixed with 500 µL of distilled water and 125 µL of Folin-Ciocaleu’s reagent. After mixing, 1250 µL of 7.0% aqueous sodium bicarbonate and 1.0 mL of distilled water were added, and the mixture was allowed to stand for 90 min at room temperature in the dark. The absorbance was measured at 760 nm using a spectrophotometer. Total phenolic concentration is expressed as mg gallic acid equivalent/g dry weight (dw). All assays were carried out in triplicate.

#### 2.2.2. Total Flavonoid Content (TFC)

Total flavonoid concentration was determined as described by Dewanto, Wu [18] with minor modifications. Briefly, the samples (250 μL) were mixed with 75 μL of 5.0% sodium nitrite followed by 150 μL of 10% of aluminum chloride, 500 μL of 1.0 M sodium hydroxide and 775 μL of distilled water. The absorbance of the mixture was measured at 510 nm. Results were expressed as mg catechin equivalents/g dw.

#### 2.2.3. Total Condensed Tannins Content (TCTC)

Condensed tannins (proanthocyanidins) were determined according to the method of Sun, Ricardo-da-Silva [19]. 3.0 mL of 4.0% vanillin solution in methanol and 1.5 mL of concentrated HCl were added to 50 µL of diluted sample. The mixture was allowed to stand for 15 min, and the absorption was measured at 500 nm against methanol as a blank. The amount of total condensed tannins is expressed as mg equivalent catechin /g dw. All samples were analyzed in triplicate.

#### 2.2.4. Quantification of phenolic compounds by LC-ESI-MS analysis

The *A. roseum* water extract was filtered through a 0.45 μm membrane before injection into the HPLC system. LC/MS analysis was performed using a LCMS-2020 quadrupole mass spectrometer (Shimadzu, Kyoto, Japan) equipped with an electrospray ionisation source (ESI) and operated in negative ionization mode. The mass spectrometer was coupled online with an ultra-fast liquid chromatography system consisted of a LC-20AD XR binary pump system, SIL-20AC XR autosampler, CTO-20AC column oven and DGU-20A 3R degasser (Shimadzu, Kyoto, Japan). An Aquasil C18 column (Thermo Electron, Dreieich, Germany) (150 mm × 3 mm, 3 μm) preceded by an Aquasil C18 guard column (10 mm × 3 mm, 3 μm, Thermo Electron) were used for analysis. The mobile phase was composed of A (0.1% formic acid in H_2_O, v/v) and B (0.1% formic acid in methanol, v/v) with a linear gradient elution: 0-45 min, 10-100% B; 45-55 min, 100% B. Re-equilibration duration was 5 min between individual runs. The flow rate of the mobile phase was 0.4 mL/min, the column temperature was maintained at 40 °C and the injection volume was 5.0 μL. The spectra of the eluted materials were monitored in SIM (Selected Ion Monitoring) mode and processed using Shimadzu Lab Solutions LC-MS software. High-purity nitrogen was used as the nebulizer and auxiliary gas. The mass spectrometer was operated in negative ion mode with a capillary voltage of −3.5 V, a nebulizing gas flow of 1.5 l/min, a dry gas flow rate of 12 l/min, a DL (dissolving line) temperature of 250 °C, a block source temperature of 400 °C, a voltage detector of 1.2 V and the full scan spectra from 50 to 2000 Da.

### 2.3. Preparation of Aβ_42_ amyloid fibrils

The Aβ_42_ fibrils were prepared as previously described [20]. The lyophilized Aβ_42_ peptide (Bachem, Bubendorf, Switzerland) was dissolved in 100% hexafluoro-2-isopropanol (HFIP) to 1.0 mM and the solvent was evaporated. Aβ_42_ fibrils were prepared by suspending the peptide at the same concentration in 50 mM NaOH and diluting this solution in PBS to a final Aβ_42_ concentration of 25 μM. Then, the sample was centrifuged at 22,000 r.c.f. for 30 min, the pellet discarded and the supernatant incubated at 25 °C without agitation for 72 h.

### 2.4. Thioflavine T assay (ThT)

Aβ_42_ aggregation was evaluated by the (ThT) assay as previously described with some modifications [21]. Aβ_42_ (25 μM) samples, incubated alone or with different concentrations of *A. roseum* extract, were diluted to 15 μM (monomeric peptide concentration) in 20 mM phosphate buffer, pH 7.4, at 25 °C and supplemented with a small volume of a 1.0 mM Thioflavin T (ThT) solution adjusted to 20 μM final concentration. Then, each sample was transferred into multiple wells of a 96- well plate (200 μL/well) and ThT fluorescence was read at 485 nm, the maximum intensity of fluorescence, using a Biotek Synergy 1H plate reader; excitation wavelength was 440 nm. Buffer fluorescence was subtracted from fluorescence values of all samples. The percent inhibition of Aß_42_ aggregation was calculated using the following formula: I (%) = [(F0-F1)/F0]*100, where F0 and F1 are the florescence of Aß_42_ and Aß_42_ + Sample at 485 nm, respectively.

### 2.5. Dynamic light scattering (DLS)

Size distribution analysis of Aß_42_ in the presence or in the absence of extracts was carried out at 25 °C on 25 μM Aß_42_ samples using a Malvern Zetasizer Nano S dynamic light scattering (DLS) device (Malvern, Worcestershire, UK). Each sample was analyzed considering the refraction index and viscosity of its dispersant. A 10 mm reduced volume plastic cell was used.

### 2.6. Transmission electron microscopy (TEM) analysis

3.0 μl aliquots of 25 μM Aβ_42_ aggregates incubated for 72 h at 25 °C with or without each extract was spotted onto a Formvar and carbon-coated nickel grid and negatively stained with 1.0% (w/v) uranyl acetate. The grid was air-dried and examined using a JEM 1010 Transmission electron microscope at 80 KV excitation voltages.

### 2.7. Cell culture and MTT assay

Human neuroblastoma SH-SY5Y cells were grown in complete culture medium containing a DMEM/Ham’s nutrient mixture F-12 (1:1) supplemented with 10% fetal calf serum (FCS, Sigma-Aldrich), glutamine and antibiotics (penicillin and streptomycin) and maintained at 37 °C under 5% CO_2_. Extract protection of SH-SY5Y cells against Aβ_42_ aggregate toxicity was determined by the MTT reduction assay. SH-SY5Y cells were seeded into 96-well plates at a density of 10^4^ cells/well and allowed to attach for 24 h. 25 μM of Aβ_42_ aggregates grown in the presence or in the absence of each extract (100 μg/mL) were diluted in the culture medium and administrated to the cells at a final concentration of 2.5 μM (monomeric peptide concentration). In other experiments, the cells were pretreated for 24 h with each extract (10, 25, 50 and 100 μg/mL: final concentration) and then exposed to Aß_42_ aggregates for 24 h. The cells were also treated with each extract (10, 25, 50 and 100 μg/mL: final concentration) in the absence of aggregated material. After 24 h, the culture medium was removed and the MTT solution (0.5 mg/mL) was added to all wells and incubated in the dark at 37 °C for 4 h. At the end of the incubation, the cells were lysed using DMSO (100%) and the amount of formazan produced was determined by measuring the absorbance at 595 nm using a Microplate reader (Biorad).

### 2.8. Reactive oxygen species measurement

The intracellular levels of reactive oxygen species (ROS) were determined using the fluorescent probe 2, 7–dichlorofluorescein diacetate, acetyl ester (CM-H_2_ DCFDA; Sigma-Aldrich), a cell permeant indicator for ROS that is fluorescent upon removal of the acetate groups by intracellular esterases and subsequent oxidation. The latter can be detected by monitoring the increase in fluorescence at 538 nm. The cells were seeded as described above for the viability assay and exposed to Aβ_42_ aggregates. After 24 h, 10 μM DCFDA in DMEM without phenol red was added for 30 min; at the end of the incubation, the fluorescence values at 538 nm were measured by a Biotek Synergy 1H plate reader.

### 2.9. Cytosolic calcium levels

The cytosolic levels of free Ca^2+^ were measured using the fluorescent probe Fluo-3 acetoxymethyl ester (Fluo-3 AM; Invitrogen, Monza). Sub-confluent SH-SY5Y cells cultured on glass coverslips were incubated with 5.0 μM Fluo-3 AM at 37 °C for 10 min prior to exposure to Aß_42_ aggregates obtained after 72 h of aggregation in the absence or in the presence of each extract (100 μg/mL) for 30 min. At the end of the incubation, the cells were fixed in 2.0% buffered paraformaldehyde for 10 min. Cell fluorescence was imaged using a confocal Leica TCS SP5 scanning microscope (Leica, Mannheim, Ge) equipped with a HeNe/Ar laser source for fluorescence measurements. The observations were performed using a Leica Plan 7 Apo X63 oil immersion objective, suited with optics for DIC acquisition. Cells from five independent experiments and three areas (about 20 cells/area) per experiment were analyzed.

### 2.10. Confocal immunofluorescence

SH-SY5Y cells grown on glass coverslips were pre-treated for 24 h with the highest concentration used for the MTT assay; 100 μg/mL of ARE extract before exposure for 24 h to Aß_42_ fibrils and Aβ_42_ grown in the presence or in the absence of extract. After incubation, the cells were washed with PBS, then, GM1 at the cell surface was labeled by incubating the cells with 10 ng/mL CTX-B Alexa 488 in cold complete medium for 30 min at room temperature. Finally, the cells were fixed in 2.0% buffered paraformaldehyde for 8 min, permeabilized by a 50% acetone/50% ethanol solution for 4.0 min at room temperature, washed with PBS and blocked with PBS containing 0.5% BSA and 0.2% gelatin. Then, the cells were incubated for 1 h at room temperature with a rabbit polyclonal antibody raised against Aß_42_ (Abcam, Cambridge, UK) diluted 1:500 in the blocking solution and washed with PBS for 30 min under stirring. The immunoreaction was revealed with Alexa 568-conjugated anti-rabbit secondary antibody (Invitrogen) diluted 1:100 in PBS. Finally, the cells were washed twice in PBS and once in water to remove non-specifically bound antibodies. Cell fluorescence was imaged using a confocal Leica TCS SP5 scanning microscope (Leica, Mannheim, Germany) equipped with a HeNe/Ar laser source for fluorescence measurements. The observations were performed using a Leica Plan 7 Apo X63 oil immersion objective. FRET analysis was performed by adopting the FRET sensitized emission method as previously reported [22].

### 2.11. Statistical analysis

All values were expressed as the mean ± SD and they were analyzed by one-way analysis of variance (ANOVA) followed by Dunnett test for MTT assay and ROS production (SPSS version 20 software). A *p*-value of less than 0.05 was considered significant.

## 3. Results

### 3.1. *A. roseum* extract (ARE) is a rich source of phenolic compounds

Total phenolic, flavonoid and condensed tannin contents (TPC, TFC and TCTC) of the ethanolic extract of *A. roseum* were determined by spectrophotometric methods. TPC measured 83.54 mg gallic acid equivalent /g dw (mg GAE / g dw). TFC and TCTC were 33.41 and 4.35 mg catechin equivalent / g dw (mg CE / g dw, respectively) (Table 1). The identification and quantification of some phenolic compounds, particularly flavonoids and phenolic acids, was obtained by chromatographic analysis. 8 compounds were identified and quantified (3 phenolic acids and 5 flavonoids) (Table 2). We detected considerable concentrations of two flavonoids; kæmpferol and the glycosylated form of luteolin (luteolin-7-o-glucoside), 869 and 683 ppm respectively.

**Table 1.** Total phenolic (TPC), total flavonoid (TFC) and total condensed tannins (TCTC) contents in ethanolic extract of *A. roseum* leaves.

|  |  |
| --- | --- |
| TPC (mg GAE / g dw) | 83.54 ± 2.63 |
| TFC (mg CE / g dw) | 33.41 ± 2.25 |
| TCTC (mg CE /g dw) | 4.35 ± 0.16 |

Data are expressed as mean ± standard deviation.

**Table 2.** Quantified phenolic compounds on ARE by HPLC-ESI-MS.

| Compounds | Concentration (mg/Kg) |
| --- | --- |
| *Phenolic acids* |  |
| Quinic acid | 4.25 ± 0.57 |
| Protocatechuic acid | 15.1 ± 2.09 |
| Trans ferulic acid | 4.05 ± 0.48 |
| *Flavonoids* |  |
| Rutin | 14.5 ± 1.93 |
| Luteolin-7-o-glucoside | 683 ± 43.85 |
| Apigenin-7-o-glucoside | 39.65 ± 4.13 |
| Kæmpferol | 869 ± 55.37 |
| Apigenin | 17.55 ± 1.89 |

Data are expressed as mean ± standard deviation.

### 3.2. ARE counteracts Aβ_42_ aggregation and inhibits its polymerization

To study the effect of ARE on Aβ_42_ aggregation, we used the ThT assay on the peptide incubated with four different concentrations of the extract (10, 25, 50 and 100 µg/mL). The kinetics of Aβ_42_ aggregation in the absence or in the presence of ARE have been followed during 72 h (Fig. 1A). The ThT fluorescence emission of Aß_42_ incubated under aggregation conditions in the absence of ARE showed a typical sigmoidal curve, with a maximum peak of fluorescence after 24 h of aggregation. However, ARE dose dependently inhibited the appearance of ThT-positive species of Aβ_42_, with an almost complete suppression at 100 µg/mL. Inhibition percent of Aβ_42_ aggregation reached its maximum after 24 h and it ranged between 38.54 % and 63.61 % when Aβ_42_ were grown in the presence of 10 and 100 µg/mL of ARE (Fig. 1B). The inhibitory effect was further confirmed by DLS, showing the size distribution analysis of Aß_42._ In the presence of ARE (100 µg/mL)Aß_42_ exhibited an hydrodynamic diameter around 13 nm until 72h of aggregation, (Fig. 1C). These results were confirmed and extended by TEM analysis; in fact, typical ordered fibrillar structures were observed in the Aβ_42_ sample incubated under aggregation conditions for 72 h, whereas, amorphous structures were imaged when Aβ_42_ was incubated in the presence of ARE and the amount of amorphous structures was increased with increasing concentration of ARE. In particular, the highest concentration (100 µg/mL) of ARE almost completely suppressed amyloid fibril formation (Fig. 1D). To asses if the positive results obtained with ARE derived from total extract or its major phenolic compound, kæmpferol, we performed a morphological analysis by TEM imaging on Aβ42 aggregation in the presence of three different concentrations of this flavonol (K1= 25 µg/mLl; K2= 50 µg/mL and K3= 100 µg/mL). Fig. 2 shows that kæmpferol on Aβ_42_ aggregation behaved similarly to ARE, resulting in amorphous structures at all tested concentrations.

**Figure 1.**
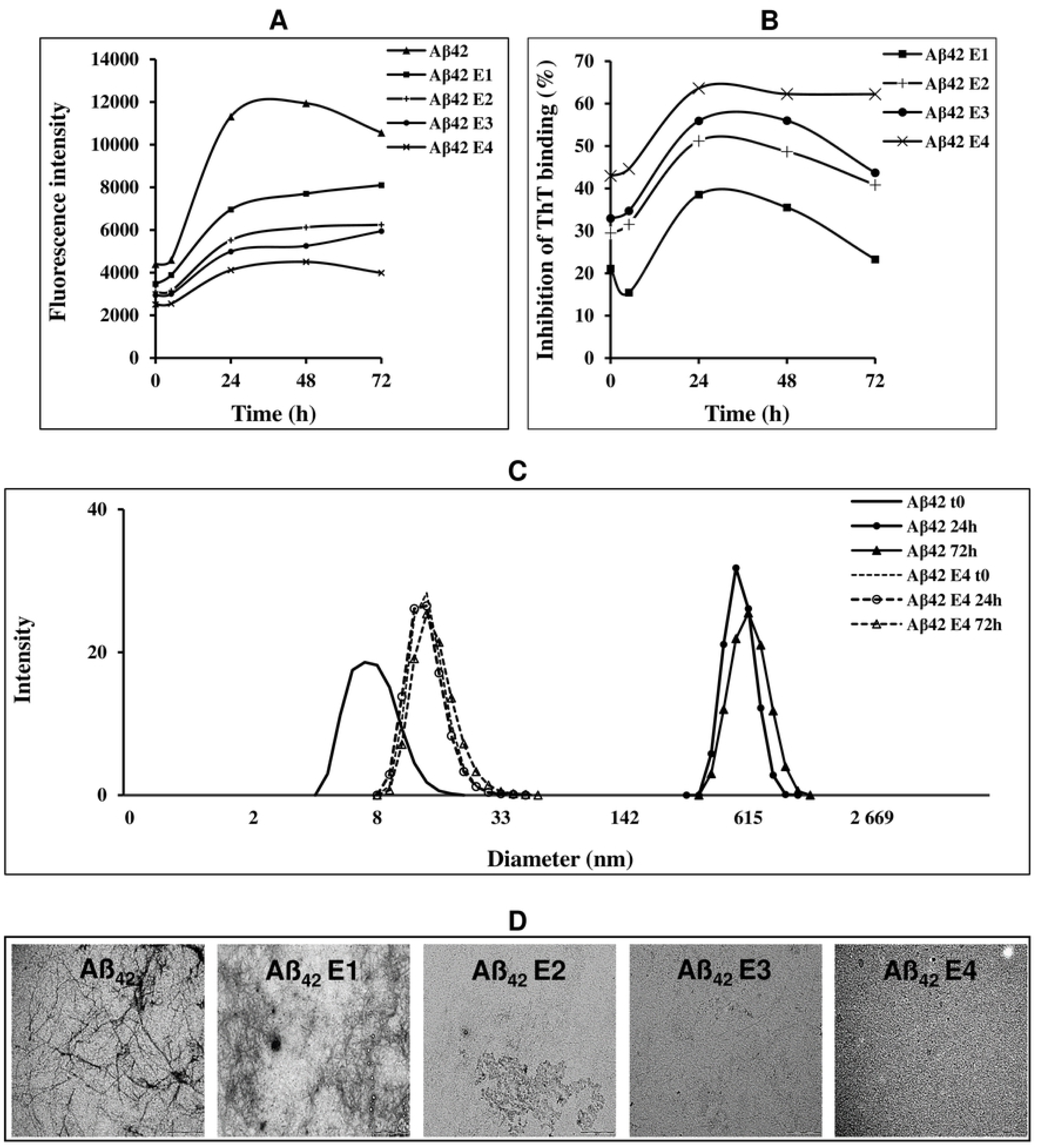
Kinetic and percent of inhibition of Aβ_42_ aggregation (A and B) and biophysical analysis of Aβ_42_ grown in the presence of different concentrations of ARE (C and D). (A). ThT fluorescence intensity in the absence and in the presence of 10, 25, 50 and 100 µg/mL (respectively E1, E2, E3 and E4) of ARE. (B). Percent inhibition of Aβ_42_ aggregation (C). DLS analysis. Size distribution of Aβ_42_ alone or aggregated in the presence of 100 µg/mL for 24 and 72 h. (D). TEM analysis. 25 μM of Aβ_42_ grown at 25 °C for 72 h in the absence or in the presence of ARE at 10, 25, 50 or 100 µg/mL.

**Figure 2.**
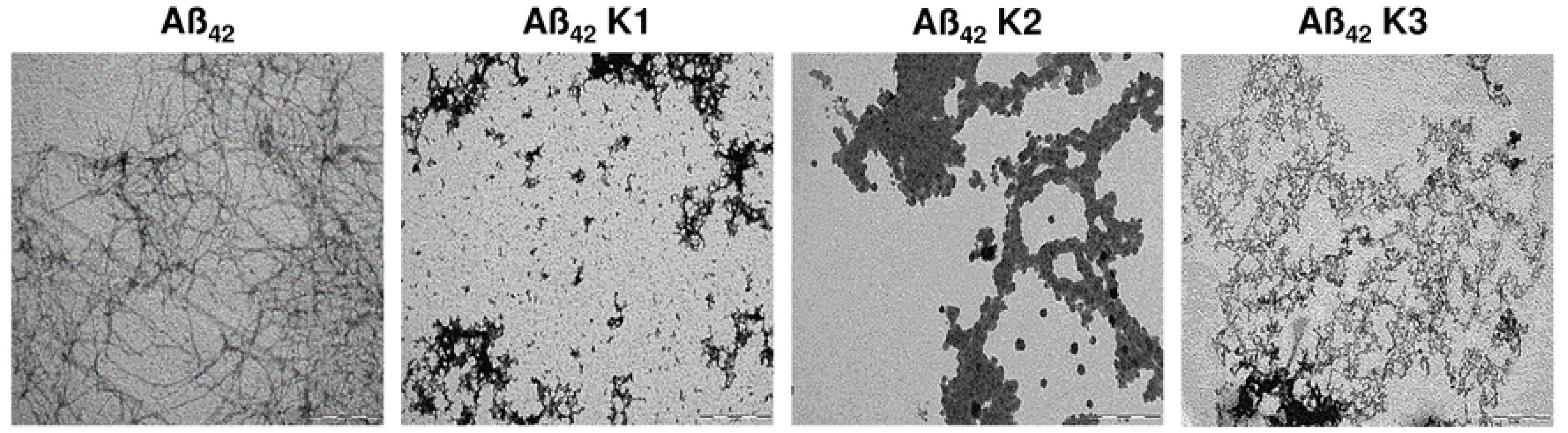
TEM analysis for kæmpferol at 25, 50 and 100 µg/mL (respectively K1, K2 and K3). 25 μM of Aβ_42_ was incubated in the presence or in the absence of each extract at 25 °C for 72 h.

Taken together, the ThT assay, DLS and TEM results show that ARE interferes with the Aβ_42_ aggregation path leading to the formation of disordered aggregates with reduced regular secondary structure in place of ordered fibrils.

### 3.3. ARE reduces Aβ_42_ cytotoxicity on SH-SY5Y neuroblastoma cells

Once characterized the biophysical features of Aβ_42_ aggregates in the absence or in the presence of various amounts of ARE or of kæmpferol, we tested the cytotoxicity of those aggregates on SH-SY5Y neuroblastoma cells by the MTT assay. We also evaluated both the possible cytotoxicity and the neuroprotective effect of ARE and kæmpferol by pre-treating the cells with either the extract or kæmpferol before adding pre-formed Aβ_42_ fibrils.

ARE did not exhibit any cytotoxic effect on SH-SY5Y neuroblastoma cells. As shown in Fig. 3A, Aβ_42_ fibrils significantly decreased cell viability; however, the cytotoxicity was strongly reduced when the cells were treated with Aβ_42_ aggregates grown in the presence of ARE. The inhibition of Aβ_42_ toxicity was dose-dependent. In fact, when Aβ_42_ was grown in the presence of 10 and 50 µg/mL of ARE, the viability of neuroblastoma cells was increased by 66.73 to 71.7 % respect to that observed in cells treated with Aβ_42_ fibrils alone. Moreover, the viability of cells exposed to Aβ_42_ aggregated in the presence of ARE 100 µg/mL was similar to that of untreated cells. These results suggest that ARE hinders Aβ_42_ fibrillation or remodels amyloid species to amorphous, disordered, aggregates devoid of toxicity. Surprisingly, we also found that ARE exhibited a neuroprotective power; in fact, the cytotoxicity of mature Aβ_42_ fibrils grown in the absence of ARE was decreased when the aggregates were administered to neuroblastoma cells pre-treated with ARE for 24 h (Fig. 3A).

**Figure 3.**
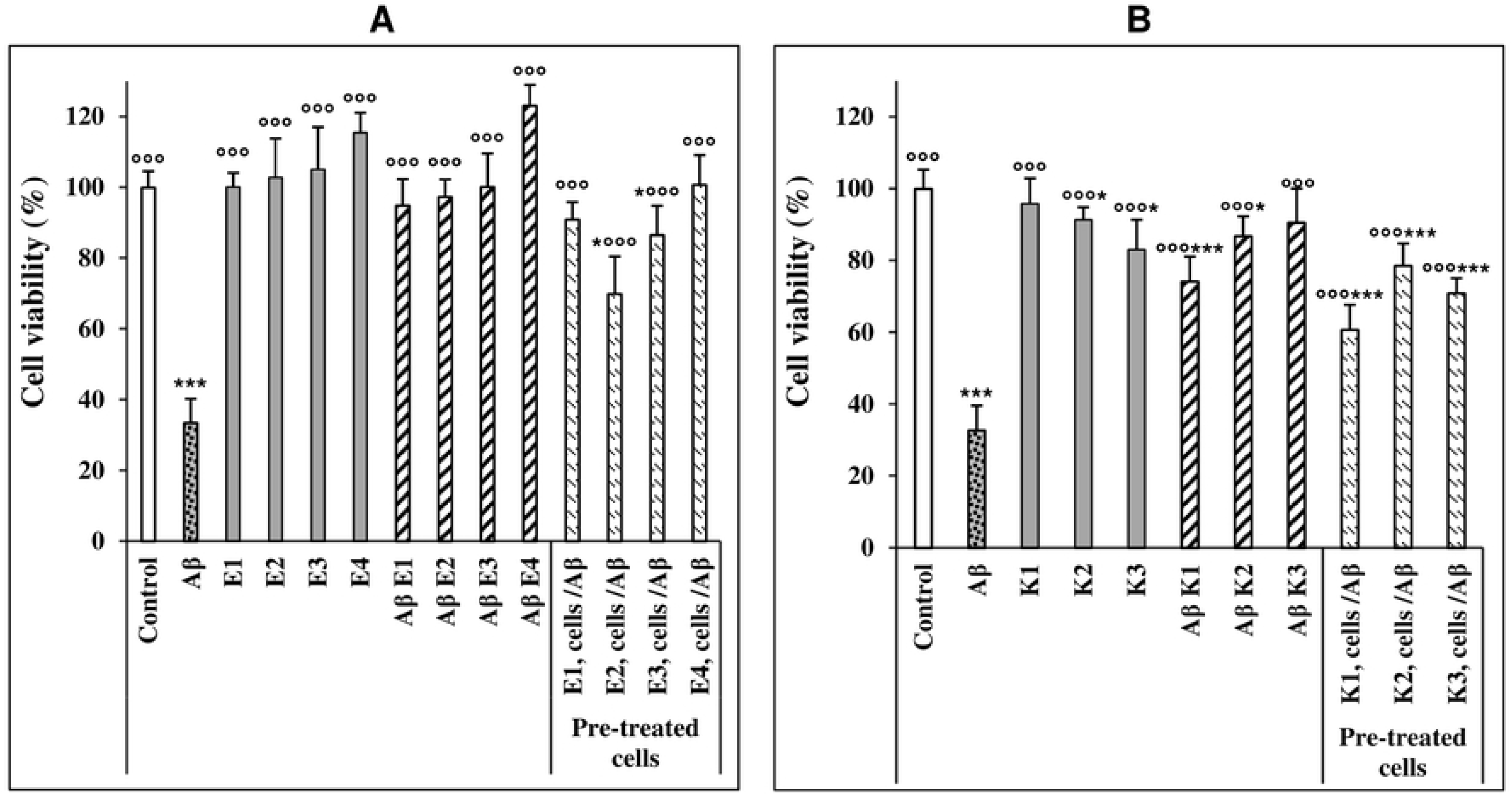
Cytotoxicity of Aβ_42_ aggregates grown in the presence or in the absence of ARE or kæmpferol. (A). MTT assay in ARE-treated cells. SH-SY5Y cells were grown in 96-well plates for 24 h then exposed to 25 μM of Aβ_42_ fibrils grown for 72 h alone or in the presence of 10, 25, 50 and 100 µg/mL ARE (E1, E2, E3 and E4, respectively). In another experiment, the cells were pretreated with the same concentrations of ARE for 24 h before exposure to 25 μM Aβ_42_ fibrils. Data are expressed as mean ± standard deviation of three independent experiments carried out in triplicate, statistical significance was performed by one-way analysis of variance (ANOVA) followed by Dunnett test; *, ** and ***or °° and °°° indicate significant statistical differences between treated and untreated control cells or versus Aβ_42_ aggregates (p < 0.05). (B) MTT assay in cells treated with kæmpferol at the same conditions described for ARE (K1= 25 µg/mL; K2 = 50 µg/mL; K3 = 100 µg/mL).

Differently from ARE, a dose dependent decrease of cell viability was seen when the cells were incubated with kæmpferol at 50 and 100 µg/mL. At these conditions, the reduction of cell viability was about 8.56 % and 16.7 %, respectively, relative to untreated cells. However, it is important to note that, similarly to ARE, kæmpferol decreased the toxicity of Aβ_42_ aggregates at all the tested concentrations such that at 100 µg/mL kæmpferol cell viability was increased from 32.72 ± 6.89 %, to 90.6 ± 9.54 % respect to untreated cells (Fig. 3B), a value approximately matching that measured in cells treated with the same kæmpferol concentration in the absence of fibrils (83.1 ± 8.32 %). We also monitored, the effects of different concentrations of kæmpferol (25, 50 and 100 µg/mL) administered to cells prior to exposure to toxic Aβ_42_ fibrils. In this case a partial reduction of fibrils toxicity was observed (from 32.72 ± 6.89 %, up 60.7 ± 6.97 %, 78.52 ± 6.31 % and 70.87 ± 4.21 %, respectively), as indicated by the MTT assay (Fig. 3B).

### 3.4. Aβ_42_ fibrils grown in the presence of ARE do not modify ROS and free calcium levels

The increase of intracellular ROS and Ca^2+^ has been investigated for the critical roles of these species in the development of oxidative stress and mitochondrial impairment involved in neurodegenerative diseases especially AD [23]. We therefore investigated (i.) the effect of Aβ_42_ aggregates grown in the presence of ARE on the intracellular ROS and Ca^2+^ levels and (ii.) the protective effect of cell pre-treatment for 24 h with different concentrations of ARE before exposure to toxic Aβ_42_ fibrils. Fig. 4A shows that, as expected, cells exposure to Aβ_42_ caused an increase of intracellular ROS up to 136.27% compared to control cells. However, a negligible ROS production was observed in cells treated with Aβ_42_ fibrils grown in the presence of different concentrations of ARE. Moreover, no apparent oxidative stress was seen in neuroblastoma cells pre-treated with different concentrations of ARE before exposure to toxic Aβ_42_ fibrils (Fig. 4A). Finally, confocal analysis showed that the intracellular level of calcium in SH-SY5Y cells exposed to Aβ_42_ aggregates grown in the presence of 100 µg/mL ARE was similar to that observed in control untreated cells whereas it decreased substantially in SH-SY5Y cells incubated with Aβ_42_ fibrils (Fig. 4B). Overall, these data suggest that ARE reduces the cytotoxicity of Aβ_42_ aggregates through their reduced ability to affect ROS and Ca^2+^ homeostasis in exposed cells possibly by affecting aggregate interaction with the cell membrane.

**Figure 4.**
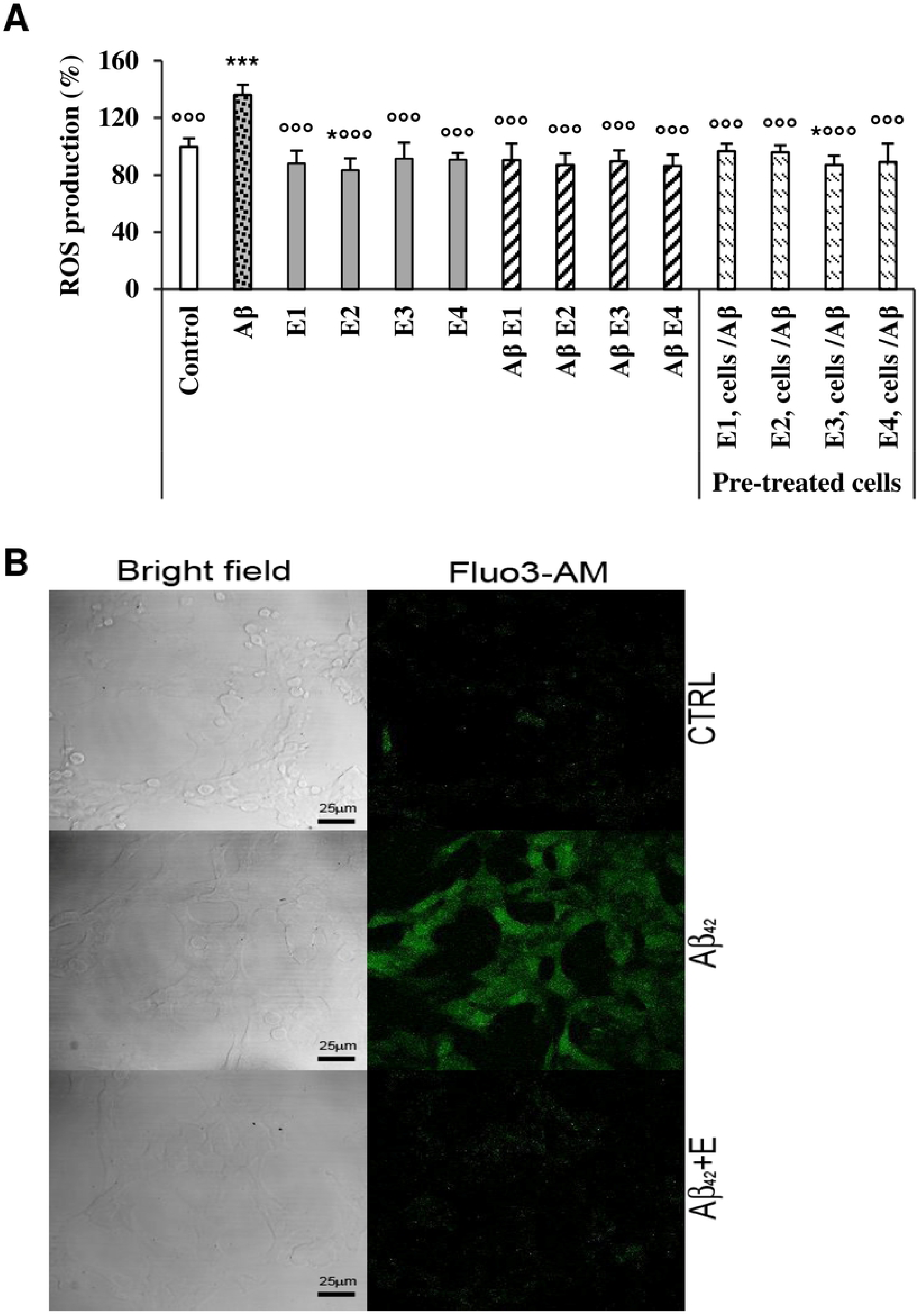
Quantification of ROS and Ca^2+^ levels. (A). Intracellular ROS production. SH-SY5Y cells were grown into 96-well plates for 24 h then exposed to 2,5 μM Aβ_42_ previously pre-incubated for 24 h alone or with 10, 25, 50 and 100 µg/mL ARE (E1, E2, E3 and E4, respectively). In another experiment, the cells were pretreated with the same concentrations or ARE for 24 h before exposure to 25 μM Aβ_42_ aggregates for 24 h. Data are expressed as mean ± standard deviation of three independent experiments carried out in triplicate, statistical significance was performed by one-way analysis of variance (ANOVA) followed by Dunnett test; * and *** or °°° indicate significant statistical differences between treated and untreated control cells or versus Aβ_42_ aggregates (p < 0.05). (B) Confocal images of free Ca^2+^ levels (green). The cells were incubated at the same conditions as in A and then treated with the fluorescent probe Fluo-3 AM for 30 min then exposed to Aβ_42_ preincubated alone or with 100 µg/mL ARE. Left: bright field. Right: Fluo3-AM fluorescence.

### 3.5. ARE reduces binding of Aβ_42_ aggregates to the cell membrane

It is widely reported that a main mechanism of amyloid aggregate cytotoxicity requires the primary interaction of the aggregates (toxic oligomers or, more frequently, mature fibrils) with the cell membrane [24], resulting in functional and/or structural perturbation of the latter. Therefore, in an additional set of experiments, we investigated the ability of the Aβ_42_ aggregates grown in the presence of ARE (100 µg/mL) to interact with the cell membrane of exposed cells with respect to the aggregates grown in the absence of ARE. To this purpose, we performed confocal microscopy experiments using a polyclonal antibody raised against recombinant Aβ_42_ and Alexa 488-conjugated CTX-B, a probe specific for the monosialotetrahexosylganglioside 1 (GM1), a common lipid raft marker widely reported as a key interaction site for amyloids [22, 25]. Aβ_42_ fibrils interaction with GM1 was evaluated by FRET analysis (Fig. 5A). As expected, a high FRET efficiency was observed in cells exposed to preformed Aβ_42_ fibrils, indicating fibrils-GM1 co-localization. However, when the cells were incubated with the same amount of Aβ_42_ fibrils grown in the presence of 100 µg/mL ARE, a reduction of both the aggregate size and aggregate amount on the cell surface was observed. Interestingly, a low interaction with toxic Aβ_42_ fibrils was elicited by the cells pretreated with the extract. The results of confocal microscopy and FRET analysis correlates with the biophysical and cytotoxicity data. Overall, these data indicate that ARE remodels Aβ_42_ fibrils to disordered species with reduced toxicity due to their inability to affect intracellular ROS and Ca^2+^ following their reduced affinity to the cell surface and impaired binding at GM1-rich sites (Fig. 5B).

**Figure 5.**
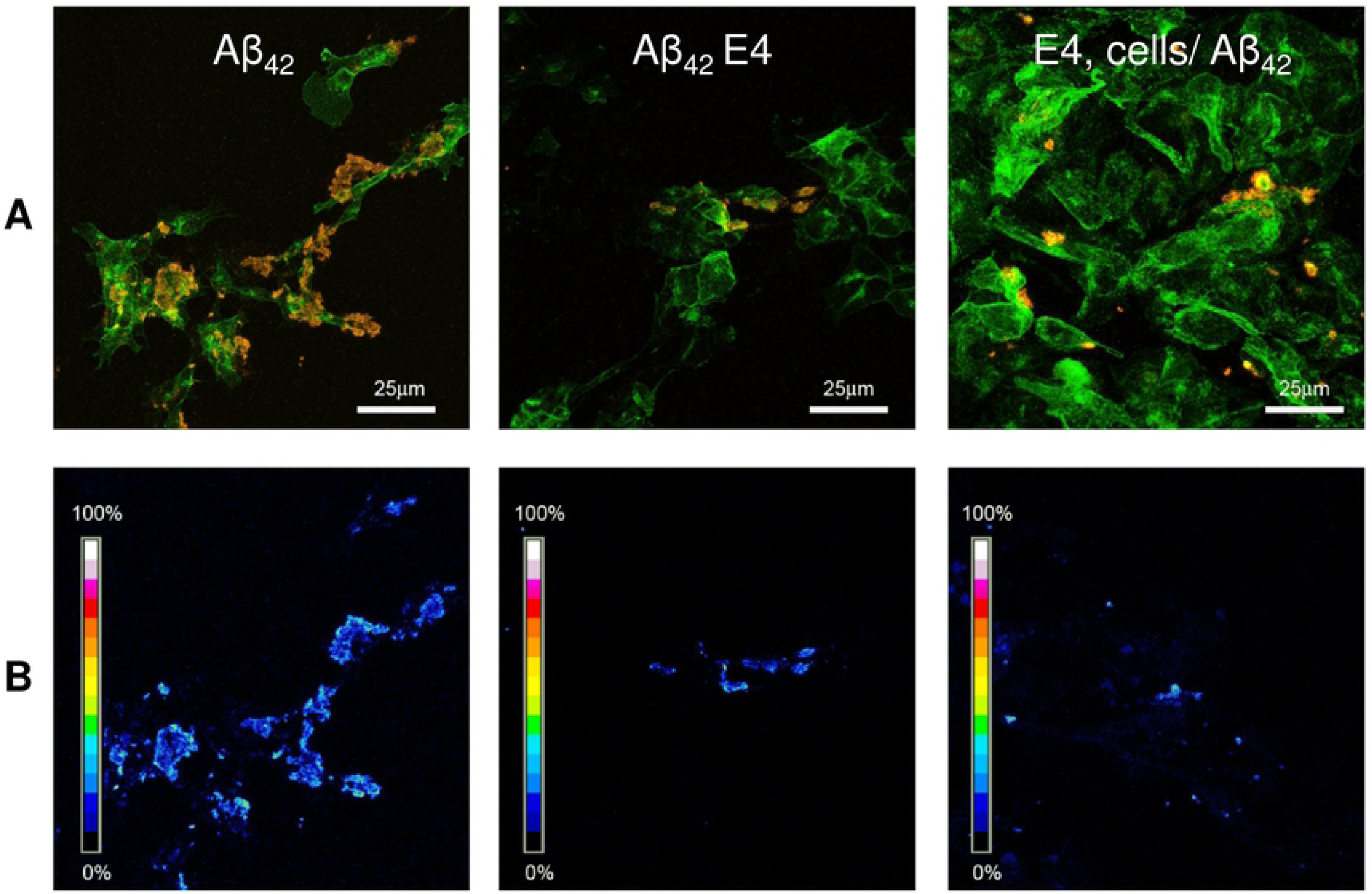
Immunolocalization of Aβ_42_ aggregates grown in the presence or in the absence of ARE. (A) Confocal imaging of SH-SY5Y treated with Aβ_42_ fibrils, Aβ_42_ fibrils grown in the presence of 100 µg/mL of ARE or the cells were pre-treated with the same concentration of ARE for 24 h before exposure to 25 μM Aβ fibrils. The cells were stained with Alexa 488-conjugated CTX-B (green fluorescence); Aβ_42_ aggregates were stained with anti-Aβ_42_ antibody followed by treatment with Alexa 568- conjugated anti-rabbit secondary antibodies (red fluorescence). (B). FRET efficiency analysis to detect the interaction between CTX-B/Alexa 488 (a GM1 marker) and anti-Aß/Alexa 568.

## 4. Discussion

The present study shows the richness in phenolic compounds of *A. roseum*. Among the eight identified molecules, kæmpferol and luteolin-7-o-glucoside were by far the most abundant compounds representing respectively 52.76% and 41.47%. In their study, Miean and Mohamed [26] reported that onion (*Allium cepa*) is the most rich plant in kæmpferol amongst 62 edible species. In the present study, we found that *A. roseum* is even richer than *A. cepa*. Furthermore, *A. roseum* is also a source of another powerful antioxidant compound, luteolin, we detected in considerable concentration as its glycosylated derivative, luteolin-7-o-glucoside. According to the same study, these two flavonoids are absent in garlic. Our results agree with those reported by Snoussi, Trabelsi [14] where the presence of five glycosylated forms of kæmpferol in *A. roseum* leaves was reported. The richness in phenolic and organosulfur compounds of *A. roseum* can be associated with its powerful antioxidant activity [14, 27], however, it still unknown if this antioxidant power matches neuroprotection.

In the present study we found that the ethanolic extract of *A. roseum* dose-dependently reduces ThT-positivity of amyloid aggregates grown from the Aβ_42_ peptide, indicating that ARE inhibits amyloid fibril formation and that this effect could be associated with the richness of this medicinal plant in phenolic compounds, particularly kæmpferol and luteolin. In their study, Sharoar, Thapa [28] found that the glycosylated form of kæmpferol, keampferol-3-o-rhamnoside, abrogates beta amyloid toxicity by modulating monomers and remodeling oligomers and fibrils to non-toxic aggregates. In another study carried out with 25 phenolic compounds, (not including kæmpferol), Churches, Caine [29] found that luteolin and transilitin display the highest inhibitory power against Aβ fibrillization. Our DLS results show that ARE also interferes with the aggregation path of Aβ_42_. The size of Aβ_42_ aggregates grown in the presence of ARE is considerably reduced respect to that of the aggregates grown in the absence of the extract. However, DLS does not distinguish between amorphous and ordered aggregates; therefore, to get more idea about the effect of ARE on the structure of aggregated Aβ_42_, we obtained TEM images of Aβ_42_ aggregated for 72 h in the absence and in the presence of four concentrations of ARE. TEM images showed the presence of amorphous aggregates even at the lowest concentration (10 µg/mL) and that the amount of these aggregates matched the increase of ARE up to the highest concentration (100 µg/mL) where all the aggregates appeared disordered. Our results are of interest, showing that this edible allium species is a rich source of molecules able to interfere with Aβ aggregation. The inhibition of Aβ fibrillization could result from the richness of ARE in kæmpferol [30]. For this reason, TEM images were also taken in the absence and in the presence of this flavonol. Similarly to ARE, kæmpferol inhibited the polymerization of Aβ_42_, suggesting that the latter affected Aβ_42_ aggregation alone, not in synergy with other molecules contained in the ARE.

In the present study, we also found that both of ARE and kæmpferol not only reduced the toxicity of Aβ_42_ aggregates to neuroblastoma cells but also protected them against the harmful effects of these dangerous aggregates. However, differently from ARE, which was non-toxic at all assayed concentrations, high concentration of kæmpferol exhibited some cytotoxicity. This finding indicates that the presence of the other molecules in ARE is important to reduce the toxic effect of kæmpferol without loss of its anti-aggregation and cell protection power. The recovery of viability of cells incubated with aggregates grown in the presence of ARE agrees with other studies indicating that protection by allium species extracts against the toxicity of Aβ_42_ aggregates [31] is associated with their richness on phenolic and organosulfur compounds. Gupta, Indi [32] found that garlic extracts possess an anti-amyloidogenic activity, which can be affected by the extraction process. Our data also show that ARE hinders Aβ_42_ aggregate binding to SH-SY5Y cell membrane at two levels: (i.) by changing their structure; (ii.) by hindering the interaction with their receptors, notably GM1, on the cell membrane, thus reducing their toxicity, as suggested by pre-treatment experiments. Indeed, the reduced interaction of the aggregates grown in the presence of ARE with the cell membrane was not merely the result of the structural changes of those aggregates; in fact, toxic Aβ_42_ fibrils added to cells pretreated with 100 µg/mL ARE were also unable to bind properly to the plasma membrane. Tsuchiya [33] reported that flavonoid compounds, including kæmpferol and luteolin, decrease membrane fluidity, confirming that these compounds can bind directly to the cell membrane, an effect likely to affect the Aβ_42_ aggregate-cell membrane interaction, with loss of cytotoxicity.

The interaction with the cell membrane is considered at the basis of amyloid aggregate cytotoxicity. In turn, the latter has been associated to the increase of intracellular Ca^2+^ and ROS, two key events in the pathogenesis of amyloidosis, particularly of neurodegenerative diseases with amyloid deposits. It has been reported that Aβ_42_ aggregates affect membrane integrity leading to an elevation of intracellular free Ca^2+^ levels [34]. The disruption of calcium homeostasis induces a cascade of events including ROS overproduction, oxidative stress and cell death mostly by apoptosis [35]. In the present study, ARE not only decreased the oxidative stress induced by ROS but also inhibited the over-production of intracellular Ca^2+^. Taken together, the results on the inhibition of aggregate-membrane binding, the reduction of the rise pf ROS/Ca^2+^ intracellular levels and those of the MTT assay, suggest that the presence of ARE in the aggregation mixture inhibits the formation of toxic Aβ_42_ aggregates possibly by interfering with aromatic stacking or even by hindering the formation of hydrophobic clusters. This hypothesis can be supported by the results obtained with other phenolic compounds active against amyloid aggregation [36]. However, we have not investigated the interference of ARE and kæmpferol with Aβ_42_ and the mechanism of its aggregation at the molecular level. Finally, it cannot be excluded that the previously reported antioxidant power of the phenolic compounds in this *Allium* extract [14, 27] could also play other protective roles. In fact, they could inhibit Aß_42_ aggregation by protecting some amino acids, especially Tyr10 and His13, against oxidation [37], a modification important for Aß_42_ self-assembly. Finally, these antioxidant compounds could also protect directly the cells against oxidative stress.

## 5. Conclusions

In conclusion, the present study indicates that ARE and its major compound kæmpferol reduce the toxicity of Aß_42_ aggregates. The ThT results show that this extract inhibits the aggregation process. The DLS data and TEM imaging indicate that it hinders the assembly of amyloid aggregates and the formation of mature fibrils. The MTT assay shows that both ARE and kæmpferol reduce and prevent Aß_42_ cytotoxicity to human neuroblastoma cells, SH-SY5Y. The immunolocalization of amyloid aggregates indicates that ARE inhibits their binding to the cell membrane. Finally, the *A. roseum* extract protects aggregate-exposed cells by counteracting the oxidative stress following reduction of ROS production and free intracellular Ca^2+^ levels.

## Acknowledgments

This work has been supported by a fellowship grant to Abdelbasset BOUBAKRI, by recurrent funding from the Tunisian Ministry of Higher Education and Scientific Research.

## Conflicts of interest

The authors declare that there are no conflicts of interest.

